# Modern machine learning outperforms GLMs at predicting spikes

**DOI:** 10.1101/111450

**Authors:** Ari S. Benjamin, Hugo L. Fernandes, Tucker Tomlinson, Pavan Ramkumar, Chris VerSteeg, Raeed Chowdhury, Lee Miller, Konrad Paul Kording

## Abstract

Neuroscience has long focused on finding encoding models that effectively ask “what predicts neural spiking?” and generalized linear models (GLMs) are a typical approach. It is often unknown how much of explainable neural activity is captured, or missed, when fitting a GLM. Here we compared the predictive performance of GLMs to three leading machine learning methods: feedforward neural networks, gradient boosted trees (using XGBoost), and stacked ensembles that combine the predictions of several methods. We predicted spike counts in macaque motor (M1) and somatosensory (S1) cortices from standard representations of reaching kinematics, and in rat hippocampal cells from open field location and orientation. In general, the modern methods (particularly XGBoost and the ensemble) produced more accurate spike predictions and were less sensitive to the preprocessing of features. This discrepancy in performance suggests that standard feature sets may often relate to neural activity in a nonlinear manner not captured by GLMs. Encoding models built with machine learning techniques, which can be largely automated, more accurately predict spikes and can offer meaningful benchmarks for simpler models.

## Introduction

A central tool of neuroscience is the tuning curve, which maps aspects of external stimuli to neural responses. The tuning curve can be used to determine what information a neuron encodes in its spikes. For a tuning curve to be meaningful it is important that it accurately predicts the neural response. Often, however, methods are chosen that sacrifice accuracy for simplicity. Since inaccurate methods may systematically miss aspects of the neural response, any choice of predictive method should be compared with methods that predict as accurately as possible.

A common predictive model for neural data is the Generalized Linear Model (GLM) (1-4). The GLM performs a nonlinear operation upon a linear combination of the input features, which are often called external covariates. Typical covariates are stimulus features, movement vectors, or the animal’s location, and may include covariate history or spike history. In the absence of history terms, the GLM is also referred to as a linear-nonlinear Poisson (LN) cascade. The nonlinear operation is usually held fixed, though it can be learned (5, 6), and the linear weights of the combined inputs are chosen to maximize the agreement between the model fit and the neural recordings. This optimization problem of weight selection is convex, allowing a global optimum, and can be solved with efficient algorithms (7). The assumption of Poisson firing statistics can often be loosened (8), as well, allowing the modeling of a broad range of neural responses. Due to its ease of use, perceived interpretability, and flexibility, the GLM has become a popular model of neural spiking.

The GLM’s central assumption is that the neural function is linear in the space of inputs (i.e. prior to the nonlinearity). The GLM thus cannot learn arbitrary multi-dimensional functions of the inputs. While this assumption may hold in certain cases (8, 9), neural responses can in general be very nonlinear (5, 10). When the nonlinearity is such that it cannot be captured by a scalar link function, a GLM will poorly predict neural activity and will learn misleading feature weights (as the optimal weight on one input will depend on the values of other inputs). To avoid this situation it is common to mathematically transform the features and thus obtain a new set that yields better spike predictions and thus better matches the linearity assumption of the GLM. In keeping with the machine learning literature, we call this step of choosing inputs *feature engineering*. When features have been found that are exactly linear with respect to neural activity after the link function, a GLM trained upon them will match or outperform more nonlinear methods. A GLM should thus be compared with benchmark methods (rather than iterating on features based on GLM fit quality alone). To check if guessed features are indeed linear with respect to neural activity prior to the nonlinearity, it is important that GLMs be compared with general nonlinear models that can express more complex stimulus–response relationships.

Machine learning (ML) methods for regression have improved dramatically since the invention of the GLM. Many ML methods require little feature engineering and do not need to assume linearity. Top performing methods, (as judged by the frequency of winning solutions on Kaggle, a ML competition website (11)) include neural networks (12), gradient boosted trees (13), and ensemble techniques. These methods are now relatively easy to implement in a few lines of code in a scripting language such as Python, enabled by well-supported machine learning packages, such as scikit-learn (14), Keras (15), Theano (16), and XGBoost (13). The greatly increased predictive power of modern ML methods is now very accessible and could improve the state of the art in encoding models across neuroscience.

Here we applied several ML methods, including artificial neural networks, gradient boosted trees, and ensembles to the task of spike prediction, and evaluated their performance alongside a GLM. We compared the methods on data from three separate brain areas. These areas differed greatly in the effect size of covariates and in their typical spike rates, and thus served to evaluate the strengths of these methods across different conditions. In each area we found that the tested ML methods could more accurately predict spiking than the GLM with typical feature choices, demonstrating both the power of these methods and the extent of nonlinearity upon the inputs that the GLM could not account for. Tuning curves built for these features with a GLM would thus not capture the full nature of neural activity. We provide our implementing code in an accessible format so that all neuroscientists may easily test and compare these methods on other datasets.

## Materials and Methods

### Data

We tested our methods at predicting spikes for neurons in the macaque primary motor cortex, the macaque primary somatosensory cortex, and the rat hippocampus. All animal use procedures were approved by the institutional animal care and use committees at Northwestern University and conform to the principles outlined in the Guide for the Care and Use of Laboratory Animals (National Institutes of Health publication no. 86-23, revised 1985). Data presented here were previously recorded for use with multiple analyses. Procedures were designed to minimize animal suffering and reduce the number used.

The macaque motor cortex data consisted of previously published electrophysiological recordings from 82 neurons in the primary motor cortex (M1) (17). The neurons were sorted from recordings made during a two-dimensional center-out reaching task with eight targets. In this task the monkey grasped the handle of a planar manipulandum that controlled a cursor on a computer screen and simultaneously measured the hand location and velocity (Fig. 1). After training, an electrode array was implanted in the arm area of area 4 on the precentral gyrus. Spikes were discriminated using offline sorter (Plexon, Inc), counted and collected in 50-ms bins. The neural recordings used here were taken in a single session lasting around 13 minutes.

**Fig 1:**
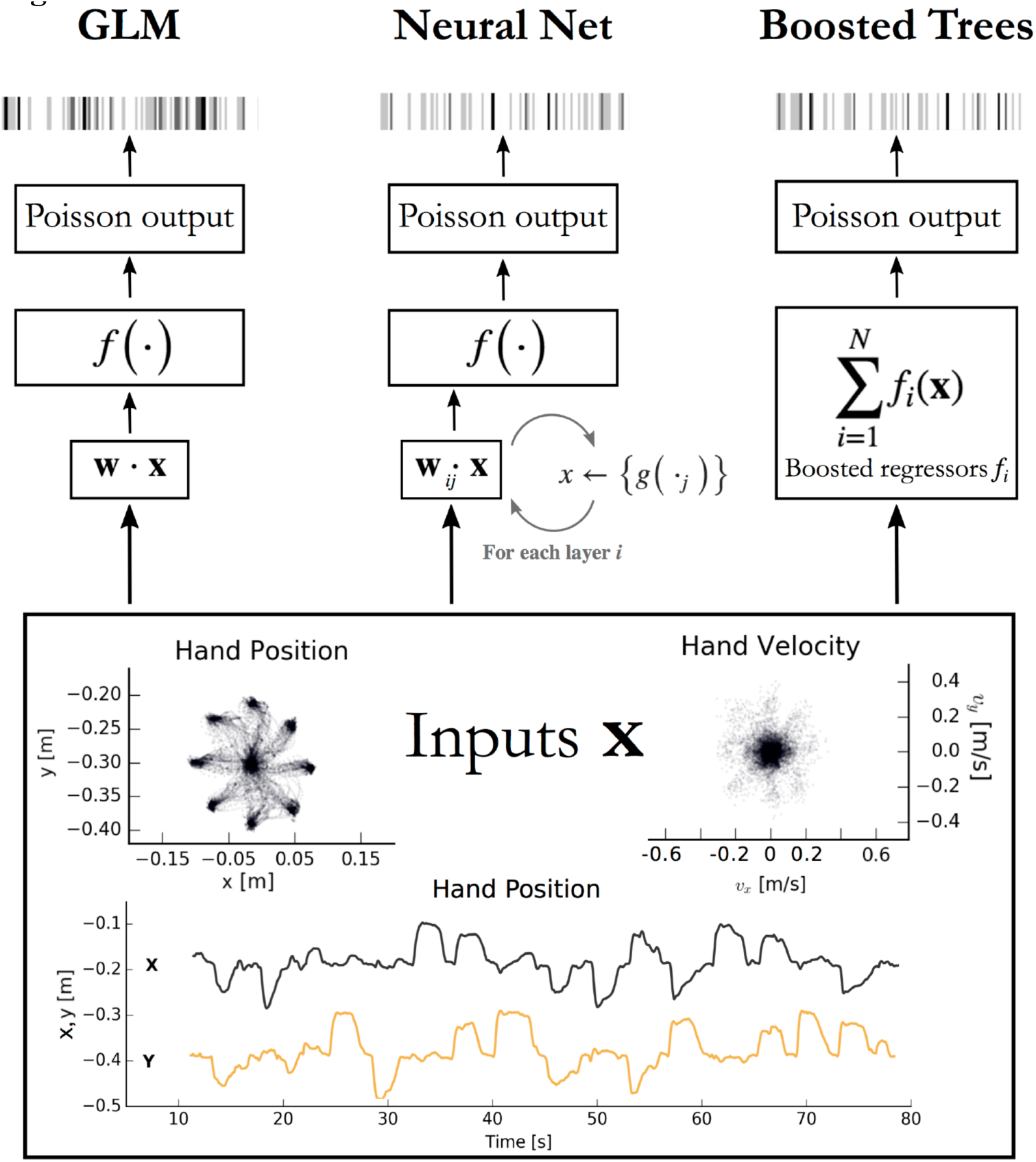
Encoding models aim to predict spikes, top, from input data, bottom. The inputs displayed are the position and velocity signals from the M1 dataset (17) but could represent any set of external covariates. The GLM takes a linear combination of the inputs, applies an exponential function *f*, and produces a Poisson spike probability that can be used to generate spikes (left). The feedforward neural network (center) does the same when the number of hidden layers *i* = 0. With *i ≥* 1 hidden layers, the process repeats; each of the *j* nodes in layer *i* computes a nonlinear function *g* of a linear combination of the previous layer. The vector of outputs from all *j* nodes is then fed as input to the nodes in the next layer, or to the final exponential *f* on the final iteration. Boosted trees (right) return the sum of N functions of the original inputs. Each of the *f*_*i*_ is built to minimize the residual error of the sum of the previous *f* _*0:i-1*_.

The macaque primary somatosensory cortex (S1) data was recorded during a two-dimensional random-pursuit reaching task and was previously unpublished. In this task, the monkey gripped the handle of the same manipulandum. The monkey was rewarded for bringing the cursor to a series of randomly positioned targets appearing on the screen. After training, an electrode array was implanted in the arm area of area 2 on the postcentral gyrus, which receives a mix of cutaneous and proprioceptive afferents. Spikes were processed as for M1. The data used for this publication derives from a single recording session lasting 51 minutes.

As with M1 (described in results), we processed the hand position, velocity, and acceleration accompanying the S1 recordings in an attempt to obtain linearized features. The features 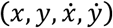 were found to be the most successful for the GLM. Since cells in the arm area of S1 have been shown to have approximately sinusoidal tuning curves relating to movement direction (18), we also tested the same feature transformations as were performed for M1 but did not observe any increase in predictive power.

The third dataset consists of recordings from 58 neurons in the CA1 region of the rat dorsal hippocampus during a single 93 minute free foraging experiment, previously published and made available online by the authors (19, 20). Position data from two head-mounted LEDs provided position and heading direction inputs. Here we binned inputs and spikes from this experiment into 50ms bins. Since many neurons in the dorsal hippocampus are responsive to the location of the rat, we processed the 2D position data into a list of squared distances from a 5x5 grid of place fields that tile the workspace. Each position feature thus has the form

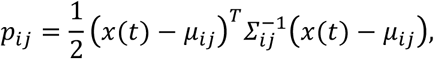

where *μ*_*ij*_ is the center of place field *i, j ≤ 5* and ∑_*ij*_ is a covariance matrix chosen for the uniformity of tiling. An exponentiated linear combination of the *p*_*ij*_ (as is performed in the GLM) evaluates to a single Gaussian centered anywhere between the place fields. The inclusion of the *p*_*ij*_ as features thus transforms the standard representation of cell-specific place fields (21) into the mathematical formulation of a GLM. The final set of features included the *p*_*ij*_ as well as the rat speed and head orientation.

### Treatment of Spike and Covariate History

We slightly modified our data preparation methods for spike prediction when spike and covariate history terms included as regressors (Fig 6). To construct spike and covariate history filters, we convolved 10 raised cosine bases (built as in (22)) with binned spikes and covariates. The longest temporal basis included times up to 250 ms before the time bin being predicted. This process resulted in 120 total covariates per sample (10 current covariates, 100 covariate temporal filters, and 10 spike history filters). We predicted spikes in 5 ms bins (rather than 50 ms) to allow for modeling of more precise time-dependent phenomena, such as refractory effects. The cross-validation scheme also differs from the main analysis of this paper, as using randomly selected splits of the data would result in the appearance in the test set of samples that were in history terms of training sets, potentially resulting in overfitting. We thus employed a cross-validation routine to split the data continuously in time, assuring that no test set sample has appeared in any form in training sets.

### Generalized Linear Model

The Poisson generalized linear model is a multivariate regression model that describes the instantaneous firing rate as a nonlinear function of a linear combination of input features (see e.g. (23, 24) for review, (22, 25, 26) for usage). Here, we took the form of the nonlinearity to be exponential, as is common in previous applications of GLMs to similar data (27). We approximate neural activity as a Poisson process, in which the probability of firing in any instant is independent of firing history. The general form of the GLM is depicted Figure 1. We implemented the GLM using elastic-net regularization, using the open-source Python package pyglmnet (28). The regularization path was optimized separately on a single neuron in each dataset on a validation set not used for scoring.

### Neural Network

Neural networks are well-known for their success at supervised learning tasks. More comprehensive reviews can be found elsewhere (12). Here, we implemented a simple feedforward neural network and, for the analysis with history terms, an LSTM, a recurrent neural network architecture that allows the modeling of time dependencies on multiple time-scales (29).

We point out that a feedforward neural network with no hidden layers is equivalent in mathematical form to a GLM (Fig. 1). For multilayer networks, one can write each hidden layer of *n* nodes as simply *n* GLMs, each taking the output of the previous layer as inputs (noting that the weights of each are chosen to maximize only the final objective function, and that the intermediate nonlinearities need not be the same as the output nonlinearity). A feedforward neural network is thus a generalization, or repeated application of a GLM.

The networks were implemented with the open-source neural network library Keras, running Theano as the backend (15, 16). The feedforward network contained two hidden layers, dense connections, rectified linear activation, and a final exponentiation. To help avoid overfitting, we allowed dropout on the first layer, included batch normalization, and allowed elastic-net regularization upon the weights (but not the bias term) of the network (30). The networks were trained to maximize the Poisson likelihood of the neural response. We optimized over the number of nodes in the first and second hidden layers, the dropout rate, and the regularization parameters for the feedforward neural network, and for the number of epochs, units, dropout rate, and batch size for the LSTM. Optimization was performed on only a subset of the data from a single neuron in each dataset, using Bayesian optimization (31) in an open-source Python implementation (32).

### Gradient Boosted Trees

A popular method in many machine learning competitions is that of gradient boosted trees. Here we describe the general operation of XGBoost, an open-source implementation that is efficient and highly scalable, works on sparse data, and easy to implement out-of-the-box (13).

XGBoost trains many sequential models to minimize the residual error of the sum of previous model. Each model is a decision tree, or more specifically a classification and regression tree (CART) (33). Training a decision tree amounts to determining a series of rule-based splits on the input to classify output. The CART algorithm generalizes this to regression by taking continuously-valued weights on each of the leaves of the decision tree.

For any predictive model 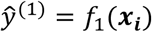 and true response *y*_*i*_, we can define a loss function 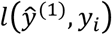 between the prediction and the response. The objective to be minimized during training is then simply the sum of the loss over each training example *i,* plus some regularizing function Ω that biases towards simple models.

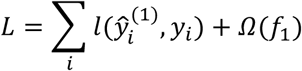

After minimizing *L* for a single tree, XGBoost constructs a second tree *f*_2_(*x*_*i*_) that approximates the residual. The objective to be minimized is thus the total loss *L* between the true response *y*_*i*_ and the sum of the predictions given by the first tree and the one to be trained.

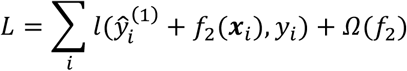

This process is continued sequentially for a predetermined number of trees, each trained to approximate the residual of the sum of previous trees. In this manner XGBoost is designed to progressively decrease the total loss with each additional tree. At the end of training, new predictions are given by the sum of the outputs of all trees.

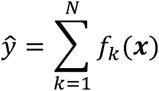

In practice, it is simpler to choose the functions *f*_*k*_ via gradient boosting, which minimizes a second order approximation of the loss function (34).

XGBoost offers several additional parameters to optimize performance and prevent overfitting. Many of these describe the training criteria for each tree. We optimized some of these parameters for a single neuron in each dataset using Bayesian optimization (again over a validation set different from the final test set). These parameters included the number of trees to train, the maximum depth of each decision tree, and the minimum weight allowed on each decision leaf, the data subsampling ratio, and the minimum gain required to create a new decision branch.

### Random Forests

We implement random forests here to increase the power of the ensemble (see below); their performance alone is displayed in Supplementary Figure 1. Random forests train multiple parallel decision trees on the features-to-spikes regression problem (not sequentially on the remaining residual, as in XGBoost) and averages their outputs (35). The variance on each decision tree is increased by training on a sample of the data drawn with replacement (i.e., bootstrapped inputs) and by choosing new splits using only a random subset of the available features. Random forests are implemented in Scikit-learn (14). It should be noted that the Scikit-learn implementation currently only minimizes the mean-squared error of the output, which is not properly applicable to Poisson processes and may cause poor performance. Despite this drawback their presence still improves the ensemble scores.

### Ensemble Method

It is a common machine learning practice to create ensembles of several trained models. Different algorithms may learn different characteristics of the data, make different types of errors, or generalize differently to new examples. Ensemble methods allow for the successes of different algorithms to be combined. Here we implemented *stacking*, in which the output of several models is taken as the input set of a new model (36). After training the GLM, neural network, random forest, and XGBoost on the features of each dataset, we trained an additional instance of XGBoost using the spike predictions of the previous methods as input. The outputs of this ‘second stage’ XGBoost are the predictions of the ensemble.

### Scoring and Cross-Validation

Each of the three methods was scored with the pseudo-R^2^ score, a scoring function applicable to Poisson processes (37). Note that a standard R^2^ score assumes Gaussian noise and cannot be applied here. The pseudo-R^2^ was calculated as

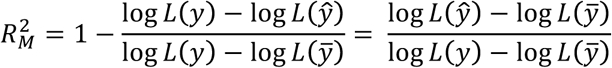

Here *L(y)* is the log likelihood of the true output, 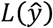 is the log likelihood of the predicted output, and 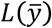 is the log likelihood, which here is the log likelihood of the data under the mean firing rate alone. The pseudo-R^2^ can be interpreted as the fraction of the maximum potential log-likelihood gain (relative to the null model) achieved by the tested model (37). The score can also be seen as one minus the ratio of the deviance of the tested model to the deviance of the null model. It takes a value of 0 when the data is as likely under the tested model as the null model, and a value of 1 when the tested model perfectly describes the data. It is empirically a lower value than a standard R2 when both are applicable (38). The null model can also be taken to be a model other than the mean firing rate (e.g. the GLM) to directly compare two methods, in which case we refer to the score as the ‘comparative pseudo-R2’. The comparative pseudo-R^2^ is referred to elsewhere as the ‘relative pseudo-R^2^’, renamed here to avoid confusion with the difference of two standard pseudo-R^2^ scores both measured against the mean (26).

We used 8-fold cross-validation (CV) when assigning a final score to the models. The input and spike data were randomly segmented, discontinuously in time, into eight equal partitions. The methods were trained on seven partitions and tested on the eighth, and this was repeated until all segments served as the test partition once. The mean of the eight scores are then recorded for the final score.

Cross-validation for ensemble methods requires extra care to ensure that there is no leak of information from the validation set into the training set. The training set for the ensemble must contain predictions from methods that were themselves not trained on the validation set. This rules out using simple *k*-fold CV with all methods trained on the same folds. Instead, we used the following nested CV scheme to train and score the ensemble. We create an outer *j=8* folds to build training and test sets for the ensemble. On each outer fold we run standard *k-*fold CV on just the training set (i.e. 7/8 of the original dataset) with each first stage method such that we obtain predictions for the whole training set of that fold. This ensures that the ensemble’s test set was never used for training any method. Finally, we build the ensemble’s test set from the predictions of the first stage methods trained on the entire training set. The process is repeated for each of the *j* folds and the mean and variance of the *j* scores of the ensemble’s predictions are recorded.

## Results

We applied several machine learning methods to predict spike counts in three brain regions and compared the quality of the predictions to those of a GLM. Our primary analysis centered on neural recordings from the macaque primary motor cortex (M1) during reaching (Fig. 1). We examined the methods’ relative performance on several sets of movement features, including one set with spike and covariate history included. Analyses of data from rhesus macaque S1 and rat hippocampus indicate how these methods compare for areas other than M1. On each of the three datasets we trained a GLM and compared it to the performance of a feedforward neural network, XGBoost (a gradient boosted trees implementation), and an ensemble method. The ensemble was an additional instance of XGBoost trained on the predictions of all three methods plus a random forest regressor. The application of these methods allowed us to demonstrate the performance of modern methods and identify whether there are typically neural nonlinearities that are not captured by a GLM. The code implementing these methods can be used by any electrophysiology lab to compare these machine learning methods with their own encoding models.

To test that all methods work reasonably well in a trivial case, we trained each to predict spiking from a simple, well-understood feature. Some neurons in M1 have been described as responding linearly to the exponentiated cosine of movement direction relative to a preferred angle (39). We therefore predicted the spiking of M1 neurons from the cosine and sine of the direction of hand movement in the reaching task. (The linear combination of a sine and cosine curve is a phase-shifted cosine, by identity, allowing the GLM to learn the proper preferred direction). We observed that each method identified a similar tuning curve (Fig. 2b) and that the bulk of the neurons in the dataset were just as well predicted by each of the methods (Fig. 2a, c) (though the ensemble was slightly more accurate than the GLM, with mean comparative pseudo-R^2^ of 0.06 [0.043 – 0.084], 95% bootstrapped confidence interval (CI)). The similar performance suggested that an exponentiated cosine is a nearly optimal approximating function of the neural response to movement direction alone, as was previously known (40). This classic example thus illustrated that all methods can in principle estimate tuning curves.

**Fig 2:**
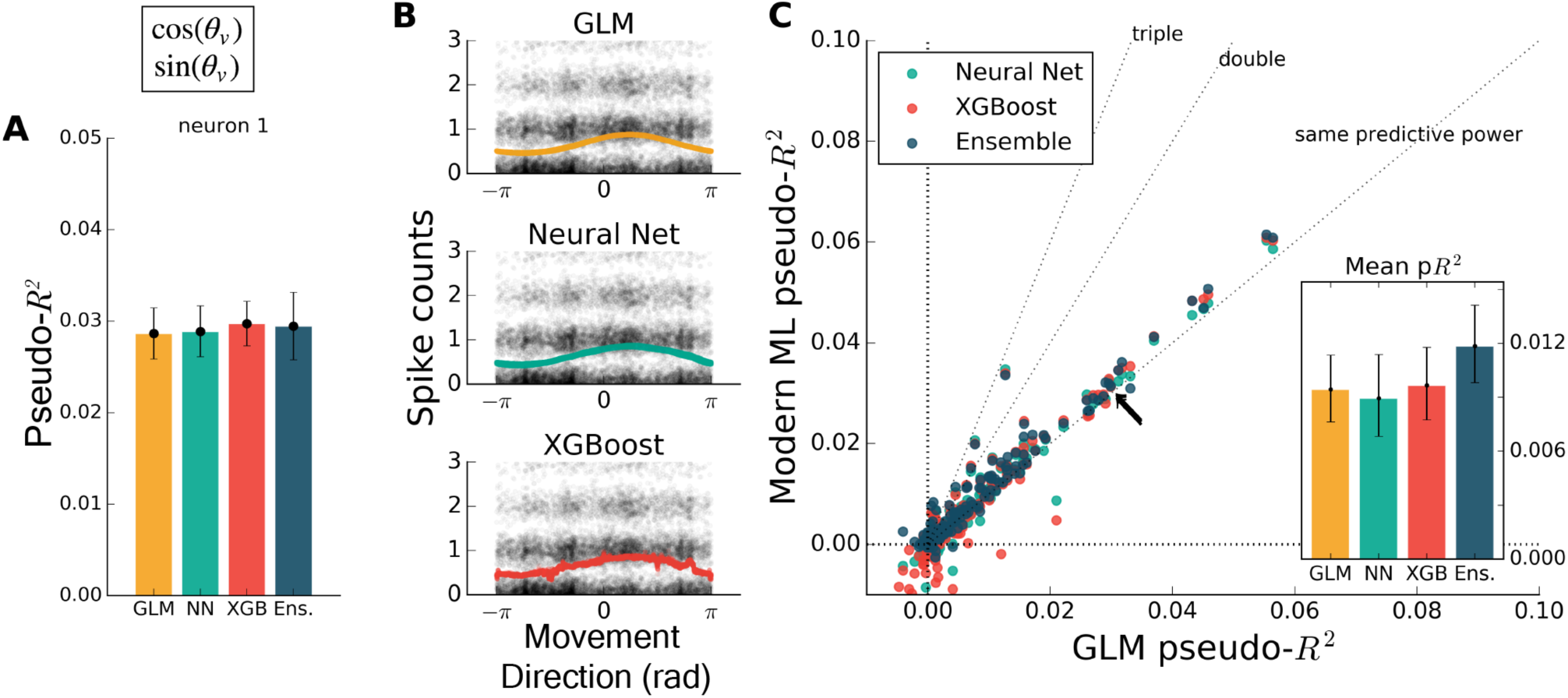
Encoding models of M1 performed similarly when trained on the sine and cosine of hand velocity direction. (a) The pseudo-R^2^ for an example neuron was similar for all four methods. On this figure and in Figures 3-5 the example neuron is the same, and is not the neuron for which method hyperparameters were optimized. (b) We constructed tuning curves by plotting the predictions of spike rate on the validation set against movement direction. The black points are the recorded responses, to which we added y-axis jitter for visualization to better show trends in the naturally quantized levels of binned spikes. The tuning curves of the neural net and XGBoost were similar to that of the GLM. The tuning curve of the ensemble method was similar and is not shown. (c) Plotting the pseudo-R^2^ of modern ML methods vs. that of the GLM indicates that the similarity of methods generalizes across neurons. The single neuron plotted at left is marked with black arrows. The mean scores, inset, indicate the overall success of the methods; error bars represent the 95% bootstrap confidence interval.

The exact form of the nonlinearity of the neural response to a given feature is rarely known prior to analysis, but this lack of knowledge need not impact our prediction ability. To illustrate the ability of modern machine learning to find the proper nonlinearity, we performed the same analysis as above but omitted the initial cosine feature-engineering step. Trained on only the hand velocity direction, in radians, which changes discontinuously at ±π, all methods but the GLM closely matched the predictive power they attained using the engineered feature (Fig. 3a). As expected, the GLM failed at generating a meaningful tuning curve (Fig. 3b). Both trends were consistent across the population of recorded neurons (Fig. 3c). The neural net, XGBoost, and ensemble methods can learn the nonlinearity of single features without requiring manual feature transformation.The inclusion of multiple features raises the possibility of nonlinear feature interactions that may elude a GLM. We found this is the case for the four-dimensional set of hand position and velocity 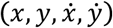. While all methods gained predictive power relative to models using movement direction alone, the GLM failed to match the other methods (Fig 4a, c). If the GLM was fit alone, and no further featuring engineering been attempted, these features would have appeared to be relatively uninformative of the neural response. If nonlinear interactions exist between preselected features, machine learning methods can potentially learn these interactions better than a GLM indicate if more linearly-related features exist.

**Fig 3:**
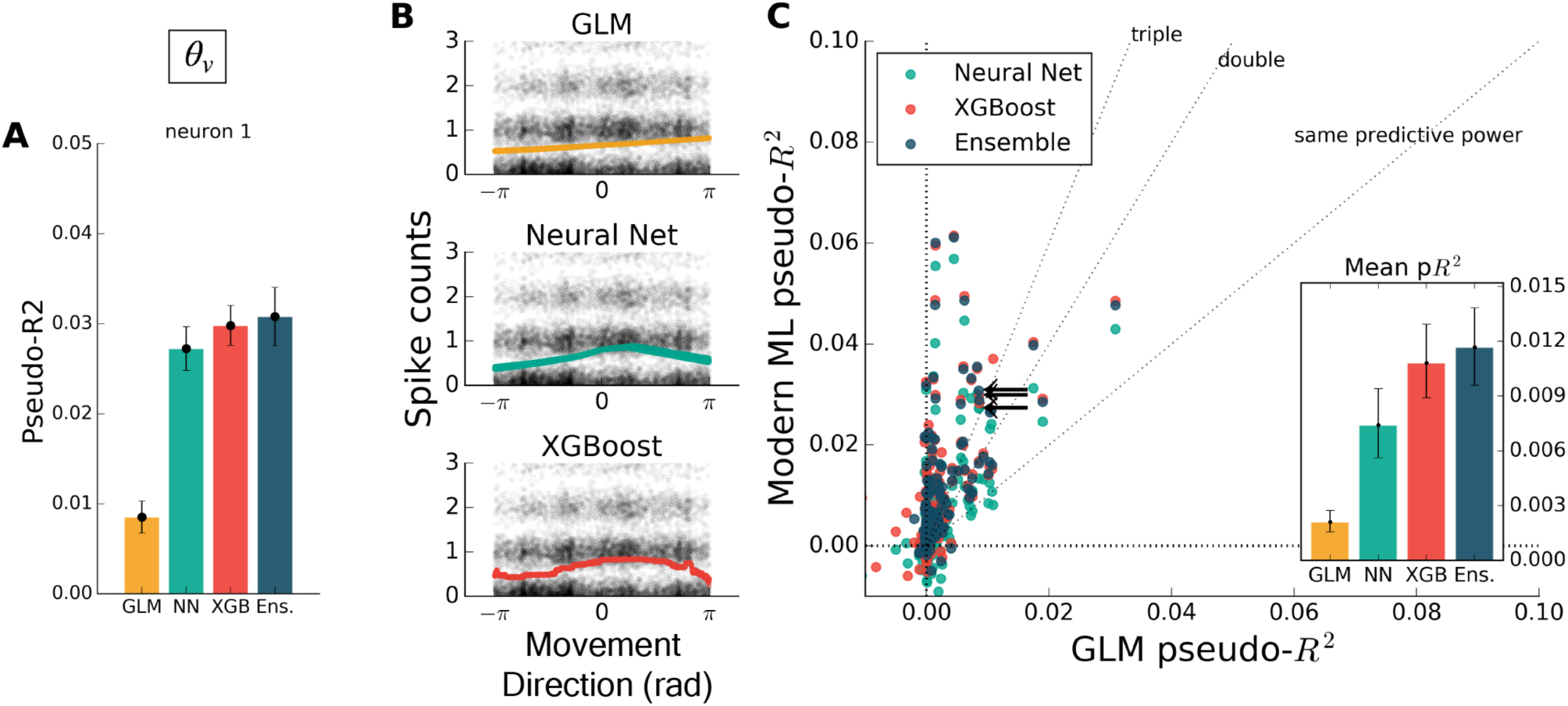
Modern ML models learn the cosine nonlinearity when trained on hand velocity direction, in radians. (a) For the same example neuron as in Figure 3, the neural net and XGBoost maintained the same predictive power, while the GLM was unable to extract a relationship between direction and spike rate. (b) XGBoost and neural nets displayed reasonable tuning curves, while the GLM reduced to the average spiking rate (with a small slope, in this case). (c) Most neurons in the population were poorly fit by the GLM, while the ML methods achieved the performance levels of Figure 2. The ensemble performed the best of the methods tested.

**Fig 4:**
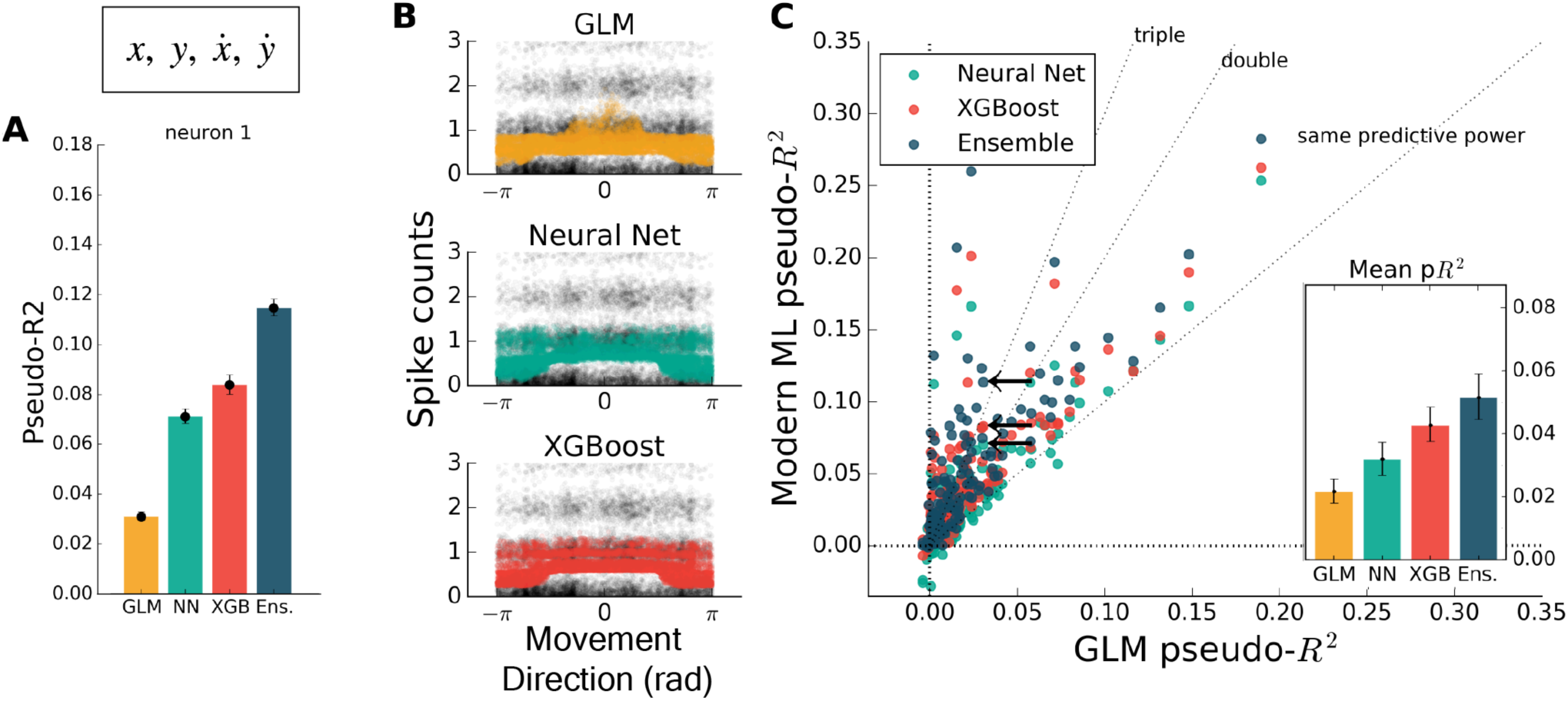
Modern ML methods can learn nonlinear interactions between features. Here the methods are trained on the feature set 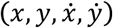. Note the change in axes scales from Figures 2-3. (a) For the same example neuron as in Figure 3, all methods gained a significant amount of predictive power, indicating a strong encoding of position and speed or their correlates. The GLM showed less predictive power than the other methods on this feature set. (b) The spike rate in black, with jitter on the y-axis, again overlaid with the predictions of the three methods plotted against velocity direction. The projection of the multidimensional tuning curve onto a 1D velocity direction dependence leaves the projected curve diffuse. (c) The ensemble method, neural network, and XGBoost performed consistently better than the GLM across the population. The mean pseudo-R2 scores show the hierarchy of success across methods.

While feature engineering can improve the performance of GLMs, it is not always simple to guess the optimal set of processed features. We demonstrated this by training all methods on features that have previously been successful at explaining spike rate in a similar center-out reaching task (6). These extra features included the sine and cosine of velocity direction (as in Figure 2), and the speed, radial distance of hand position, and the sine and cosine of position direction. The training set was thus 10-dimensional, though highly redundant, and was aimed at maximizing the predictive power of the GLM. Feature engineering improved the predictive power of all methods to variable degrees, with the GLM improving to the level of the neural network (Fig. 5). XGBoost and the ensemble still predicted spikes better than the GLM (Fig. 5c), with the ensemble scoring on average nearly double the GLM (ratio of population means of 1.8 [1.4 – 2.2], 95% bootstrapped CI). The ensemble was significantly better than XGBoost (mean comparative pseudo-R^2^ of 0.08 [0.055 – 0.103], 95% bootstrapped CI) and was thus consistently the best predictor. Though standard feature engineering greatly improved the GLM, the ensemble and XGBoost still could identify neural nonlinearity missed by the GLM.

**Fig 5:**
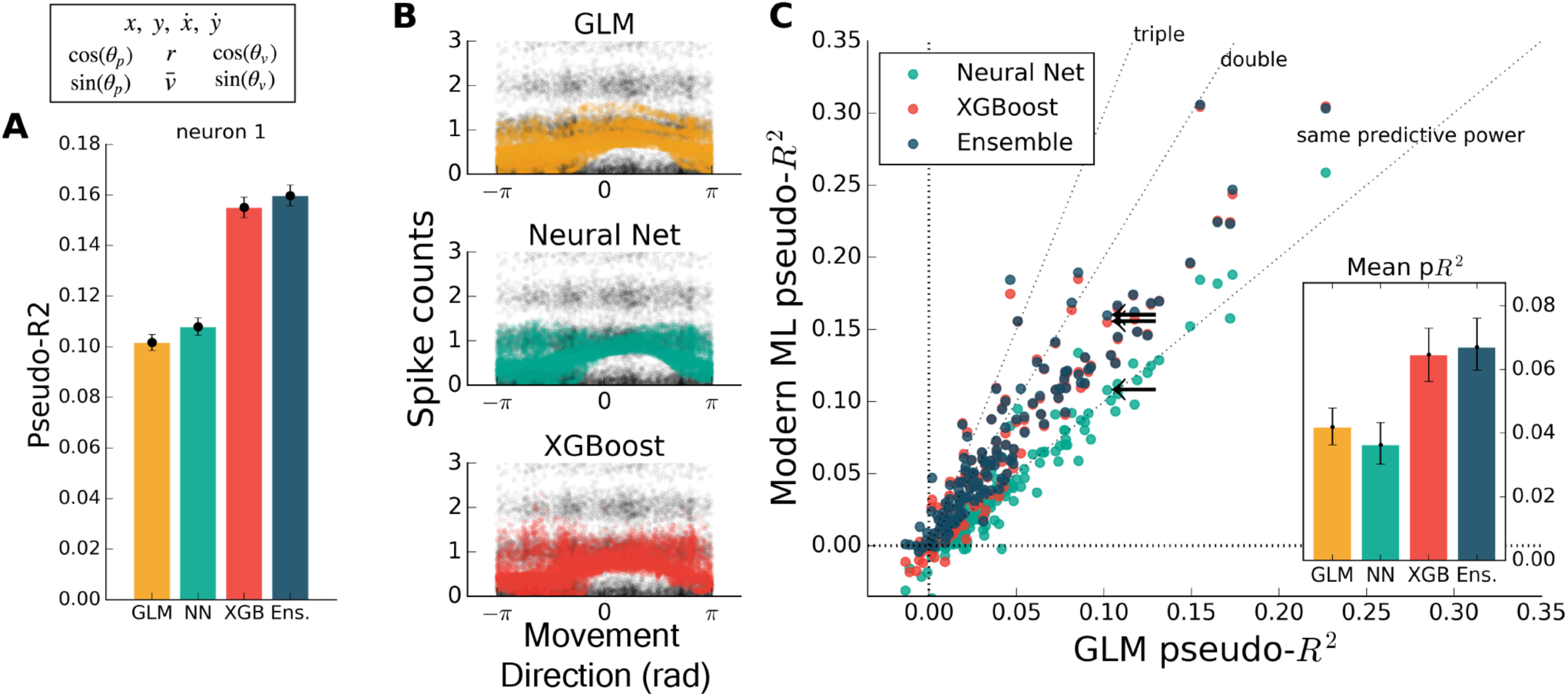
Modern ML methods can outperform the GLM even with standard featuring engineering. For this figure, all methods were trained on the features 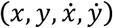 plus the engineered features. (a) For this example neuron, inclusion of the computed features increased the predictive power of the GLM to the level of the neural net. All methods increased in predictive power. (b) The tuning curves for the example neuron are diffuse when projected onto the movement direction, indicating a high-dimensional dependence. (c) Even with feature engineering, XGBoost and the ensemble consistently achieve pseudo-R^2^ scores higher than the GLM, though the neural net does not. The neuron selected at left is marked with black arrows.

**Fig 6:**
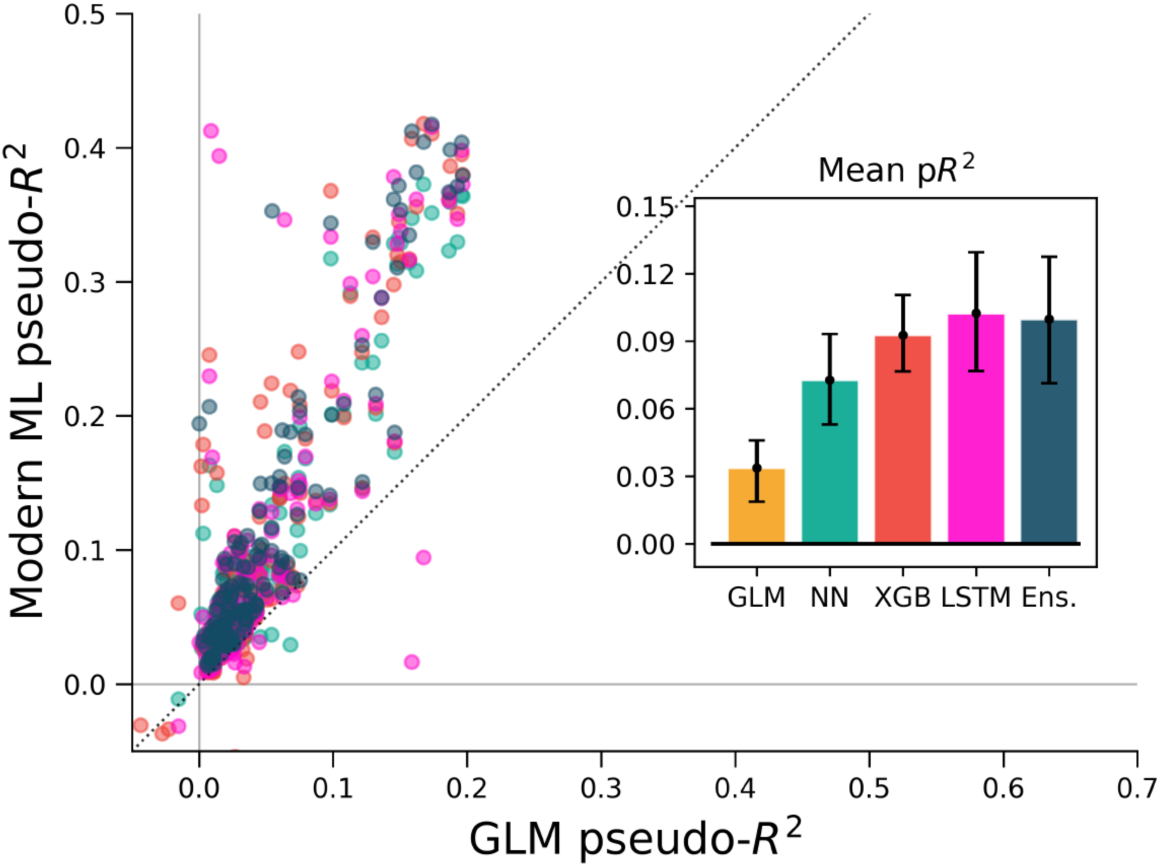
ML algorithms outperform a GLM when covariate history and neuron spike history are included. The feature set of Fig 5 (in macaque M1) was augmented with spike and covariate history terms, so that spike rate was predicted for each 5 ms time bin from the past 250 ms of covariates and neural activity. Cross-validation methods for this figure differ from other figures (see methods) and pseudo-R^2^ scores should not be compared directly across figures. All methods outperform the GLM, indicating that the inclusion of history terms does not alone allow the GLM to capture the full nonlinear relationship between covariates and spike rate.

Studies employing a GLM often include spike history as a covariate when predicting spikes, as well as past values of the covariates themselves, and it is known that this allows GLMs to model a wider range of phenomena (41). We tested select ML methods on the M1 dataset using this history-augmented feature set to see if all methods would still explain a similar level of activity. We binned data by 5 ms (rather than 50 ms) to agree in timescale with similar studies, and built temporal filters by convolving 10 raised-cosine bases with features and spikes. We note that smaller time bins result in a sparser dataset, and thus pseudo-R^2^ scores cannot be directly compared with other analysis in this paper. On this problem, our selected ML algorithms again outperformed the GLM (Fig 6). The overall best algorithm was the LSTM, which we include here as it specifically designed for modeling time series, though for most neurons XGBoost performed similarly. Thus, for M1 neurons, the GLM did not capture all predicable phenomena even when spike and covariate history were included.

To ensure that these results were not specific to the motor cortex, we extended the same analyses to primary somatosensory cortex (S1) data. We again predicted neural activity from hand movement and speed, and here without spike or covariate history terms. The ML methods outperformed the GLM for all but three of the 52 neurons, indicating that firing rates in S1 generally relate nonlinearly to hand position and velocity (Fig 7a). Each of the three ML methods performed similarly for each neuron. The S1 neural function was thus equally learnable by each method, which is surprising given the dissimilarity of the neural network and XGBoost algorithms. This situation would occur if learning has saturated near ground truth, though this cannot be proven definitively to be the case. It is at least clear from the underperformance of the GLM that the relationship of S1 activity to these covariates is nonlinear beyond the assumptions of the GLM.

**Fig 7:**
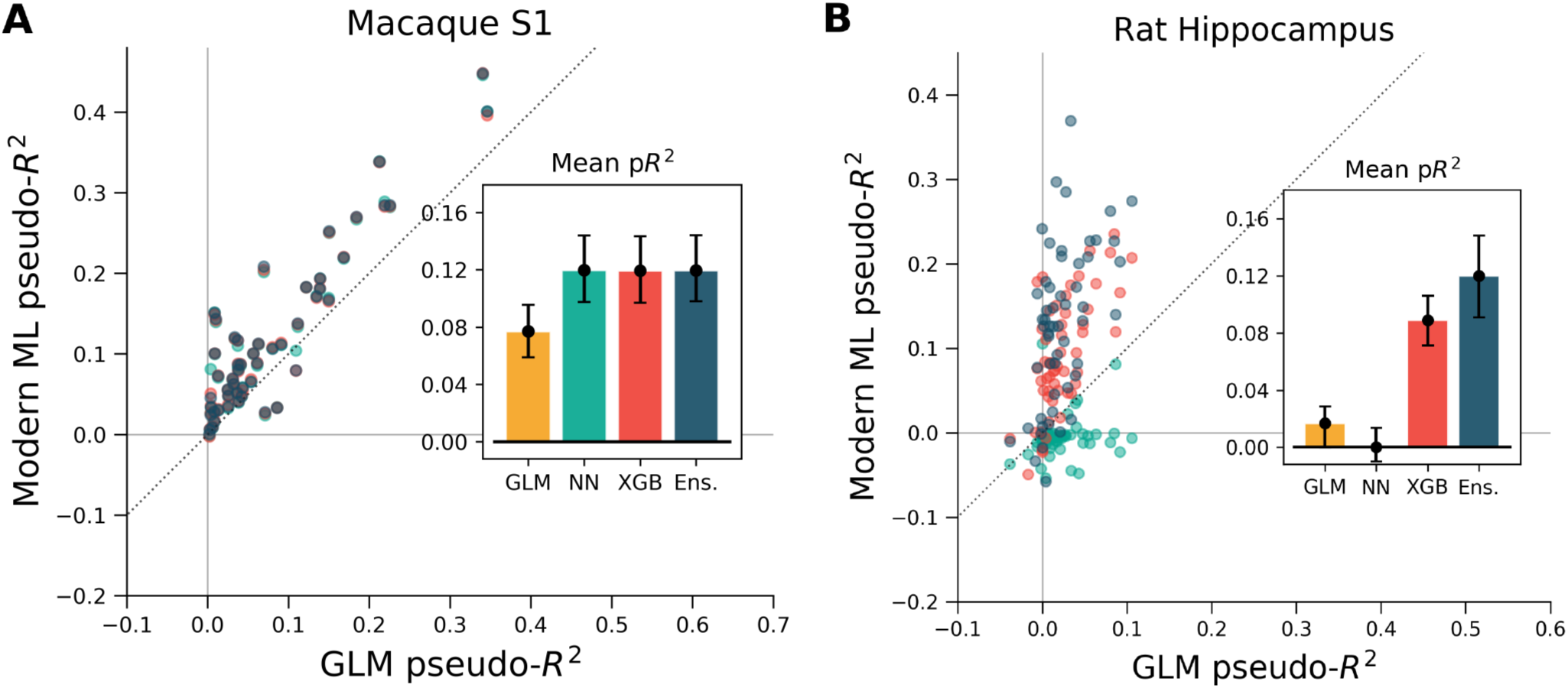
XGBoost and the ensemble method predicted the activity of neurons in S1 and in hippocampus better than a GLM. The diagonal dotted line in both plots is the line of equal predictive power with the GLM. (a) All methods outperform the GLM in the macaque S1 dataset. Interestingly, the neural network, XGBoost and the ensemble scored very similarly for each neuron in the 52 neuron dataset. (b) Many neurons in the rat hippocampus were described well by XGBoost and the ensemble but poorly by the GLM and the neural network. The poor neural network performance in the hippocampus was due to the low rate of firing of most neurons in the dataset (Supp. Fig. 2). N

We asked if the same trends of performance would hold for the rat hippocampus dataset, which was characterized by very low mean firing rates but strong effect sizes. All methods were trained on a list of squared distances to a grid of place fields and on and the rat head orientation, as described in methods. Far more even than the neocortical data, neurons were described much better by XGBoost and the ensemble method than by the GLM (Fig. 7b). Many neurons shifted from being completely unpredictable by the GLM (pseudo-R^2^ near zero) to very predictable by XGBoost and the ensemble (pseudo-R^2^ above 0.2). These neurons thus have responses that do not correlate with firing in any one Gaussian place field. We note that the neural network performed poorly, likely due to the very low firing rates of most hippocampal cells (Supp. Fig. 2). The median spike rate of the 58 neurons in the dataset was just 0.2 spikes/s, and it was only on the four neurons with rates above 1 spikes/s that the neural network achieved pseudo-R^2^ scores comparable to the GLM. The relative success of XGBoost was interesting given the failure of the neural network, and supported the general observation that boosted trees can work well with smaller and sparser datasets than those that neural networks generally require. Thus for hippocampal cells, a method leveraging decision trees such as XGBoost or the ensemble is able to capture more structure in the neural response and thus demonstrate a deficiency of the parameterization of the GLM.

## Discussion

We contrasted the performance of GLMs with various machine learning techniques at the task of predicting binned spikes in three brain regions. We found that the tested ML methods, especially XGBoost and the ensemble, routinely predicted spikes more accurately than did the GLM. Typical feature engineering only partially bridged the performance gap. Machine learning methods, especially LSTMs, also outperformed GLMs when history terms were included. The ML methods performed comparably well with and without feature engineering, even for the very low spike rates of the hippocampus dataset, indicating they could serve as convenient performance benchmarks for improving simpler encoding models. These findings indicate that instances where standard ML methods outperform GLMs may be common for neural data and standard feature choices.

When a GLM fails to explain data as well as more expressive, nonlinear methods, the current parameterization of inputs must relate to the data more nonlinearly than is assumed by the GLM. This unaccounted nonlinearity may produce feature weights that do not reflect true feature importance. A GLM will incorrectly predict no dependence on feature *x* whatsoever, for example, in the extreme case when the neural response to some feature *x* does not correlate with exp(*x*). The only way to ensure reliable interpretations of feature weights as importances is to find an input parameterization that maximizes the GLM’s predictive power. ML methods can assist this process by acting as benchmarks and indicating how much nonlinearity remains to be explained. This point relates to a possible objection to this work, which is that the GLM underperforms simply because we have selected the wrong input features. This is indeed true, as it is always theoretically possible to linearize features such that a GLM obtains equal predictive power. We mean to point out here that ML methods can highlight the deficiency of features that might have otherwise seemed uncontroversial. When applying a GLM to neural data, then, it is important to compare its predictive power with standard ML methods to ensure the neural response is properly understood.

Advanced ML methods are not widely considered to be interpretable, and some may worry that this diminishes their scientific value as encoding models. We can better discuss this issue with a more precise definition of interpretability. Following Lipton, we make the distinction between a method’s *post-hoc interpretability*, the ease of justifying its predictions, and *transparency,* the degree to which its operation and internal parameters are human-readable or easily understandable (42). A GLM is certainly more transparent than many ML methods due to its algorithmic simplicity. It is also generally more conducive to post-hoc predictions, though here is there room for modern ML methods. It is possible, for example, to visualize the aspects of stimuli that most elicit a predicted response, as has been implemented in previous applications of neural networks to spike prediction (43, 44). Various other methods exist in the literature to enable post-hoc explanations (45, 46). Here we highlight Local Interpretable Model-Agnostic Explanations (LIME), an approach that fits simple models in the vicinity of single examples to allow, at least, a local interpretation (47). On problems where interpretability is important, such capabilities for post-hoc justifications may prove sufficient.

Not all types of interpretability are necessary for a given task, and many scientific questions can be answered based on predictive ability alone. Questions of the form, “does feature *x* contribute to neural activity?”, for example, or “is past activity necessary to explain current activity?” require no method transparency. One can simply ask whether predictive power increases with feature *x*’s inclusion or decreases upon its exclusion. Importance measures based on inclusion and exclusion, or upon the strategy of shuffling a covariate of interest, are well-studied in statistics and machine learning (48, 49). Depending on the application, it may thus be worthwhile to ask not just whether different features could improve a GLM but also whether it is enough to use ML methods directly. When determining if a neuron encodes a certain set of features, that is, like muscle forces or body position, one can choose to ask if there is a neural response that is linear with respect to those features, or alternatively if there is any learnable response at all. It is possible for many questions to stay agnostic to the form of linearized features and directly use changes in predictive ability.

With ongoing progress in machine learning, many standard techniques are easy to implement and can even be automated. Ensemble methods, for example, remove the need to choose any one algorithm. Moreover, the choice of model-specific parameters is made easy by hyperparameter search methods and optimizers. In this sense, many aspects of the machine learning process are automated in our approach. We hope that this ease of use might encourage use in the neurosciences, thereby increasing the power and efficiency of studies involving neural prediction without requiring complicated, application-specific methods development (e.g. (50)). Machine learning methods are likely to be even easier to apply in the near future due to further automation. Community-supported projects in automated machine learning, such as autoSKlearn and auto-Weka, are quickly improving and promise to handle the entire regression workflow (51, 52). Applied to neuroscience, these tools will allow researchers to gain descriptive power over current methods even with simple, out-of-the-box implementations.

Machine learning methods perform quite well and make minimal assumptions about the form of neural encoding. Models that seek to understand the form of the neural code can test if they systematically misconstrue the relationship between stimulus and response by comparing their performance to these benchmarks. Encoding models built with machine learning can thus greatly aid the construction of models that capture arbitrary nonlinearity and more accurately describe neural activity.

The code used for this publication is available at https://github.com/KordingLab/spykesML. We invite researchers to adapt it freely for future problems of neural prediction.

## Author Contributions

K.K., P.R., and H.F. first conceived the project. T.T, C.V., and R.C. gathered and curated macaque data. A.B. prepared the manuscript and performed the analyses, for which H.F. and P.R. assisted. L.M. and K.K. supervised, and all authors assisted in editing.

## References

1. Simoncelli EP, Paninski L, Pillow J, Schwartz O. Characterization of neural responses with stochastic stimuli. The cognitive neurosciences. 2004;3:327–38.

2. Wu MC-K, David SV, Gallant JL. Complete functional characterization of sensory neurons by system identification. Annu Rev Neurosci. 2006;29:477–505.

3. Gerwinn S, Macke JH, Bethge M. Bayesian inference for generalized linear models for spiking neurons. Frontiers in Computational Neuroscience. 2010;4(12).

4. Nelder JA, Baker RJ. Generalized linear models. Encyclopedia of statistical sciences. 1972.

5. Chichilnisky E. A simple white noise analysis of neuronal light responses. Network: Computation in Neural Systems.2001;12(2):199–213.

6. Paninski L, Fellows MR, Hatsopoulos NG, Donoghue JP. Spatiotemporal tuning of motor cortical neurons for hand position and velocity. Journal of neurophysiology. 2004;91(1):515–32.

7. Paninski L. Maximum likelihood estimation of cascade point-process neural encoding models. Network: Computation in Neural Systems. 2004;15(4):243–62.

8. Pillow JW, Paninski L, Uzzell VJ, Simoncelli EP, Chichilnisky E. Prediction and decoding of retinal ganglion cell responses with a probabilistic spiking model. The Journal of neuroscience. 2005;25(47):11003–13.

9. Georgopoulos AP, Schwartz AB, Kettner RE. Neuronal population coding of movement direction. Science.1986;233(4771):1416–9.

10. Van Steveninck RDR, Bialek W. Real-time performance of a movement-sensitive neuron in the blowfly visual system:coding and information transfer in short spike sequences. Proceedings of the Royal Society of London B: Biological Sciences.1988;234(1277):379–414.

11. Kaggle Winner's Blog. 2016.

12. Schmidhuber J. Deep learning in neural networks: An overview. Neural Networks. 2015;61:85–117.

13. Chen T, Guestrin C. Xgboost: A scalable tree boosting system. arXiv preprint arXiv:160302754. 2016.

14. Pedregosa F, Varoquaux G, Gramfort A, Michel V, Thirion B, Grisel O, et al. Scikit-learn: Machine learning in Python. Journal of Machine Learning Research. 2011;12(Oct):2825–30.

15. Chollet F. keras. GitHub repository. 2015.

16. Team TTD, Al-Rfou R, Alain G, Almahairi A, Angermueller C, Bahdanau D, et al. Theano: A Python framework for fast computation of mathematical expressions. arXiv preprint arXiv:160502688. 2016.

17. Stevenson IH, Cherian A, London BM, Sachs NA, Lindberg E, Reimer J, et al. Statistical assessment of the stability of neural movement representations. Journal of neurophysiology. 2011;106(2):764–74.

18. Prud'homme M, Kalaska JF. Proprioceptive activity in primate primary somatosensory cortex during active arm reaching movements. Journal of neurophysiology. 1994;72(5):2280–301.

19. Mizuseki K, Sirota A, Pastalkova E, Buzsáki G. Multi-unit recordings from the rat hippocampus made during open field foraging. 2009.

20. Mizuseki K, Sirota A, Pastalkova E, Buzsáki G. Theta oscillations provide temporal windows for local circuit computation in the entorhinal-hippocampal loop. Neuron. 2009;64(2):267–80.

21. Brown EN, Frank LM, Tang D, Quirk MC, Wilson MA. A statistical paradigm for neural spike train decoding applied to position prediction from ensemble firing patterns of rat hippocampal place cells. The Journal of Neuroscience.1998;18(18):7411–25.

22. Pillow JW, Shlens J, Paninski L, Sher A, Litke AM, Chichilnisky E, et al. Spatio-temporal correlations and visual signalling in a complete neuronal population. Nature. 2008;454(7207):995–9.

23. Aljadeff J, Lansdell BJ, Fairhall AL, Kleinfeld D. Analysis of neuronal spike trains, deconstructed. Neuron. 2016;91(2):221–59.

24. Schwartz O, Pillow JW, Rust NC, Simoncelli EP. Spike-triggered neural characterization. Journal of Vision. 2006;6(4):13–25.

25. Ramkumar P, Lawlor PN, Glaser JI, Wood DK, Phillips AN, Segraves MA, et al. Feature-based attention and spatial selection in frontal eye fields during natural scene search. Journal of neurophysiology. 2016:jn. 01044.2015.

26. Fernandes HL, Stevenson IH, Phillips AN, Segraves MA, Kording KP. Saliency and saccade encoding in the frontal eye field during natural scene search. Cerebral Cortex. 2014;24(12):3232–45.

27. Saleh M, Takahashi K, Hatsopoulos NG. Encoding of coordinated reach and grasp trajectories in primary motor cortex. The Journal of Neuroscience. 2012;32(4):1220–32.

28. Ramkumar P, Jas M, Achakulvisut T, Idrizovic A, themantalope, Acuna DE, et al. Pyglmnet 1.0.1. 2017.

29. Gers FA, Schmidhuber J, Cummins F. Learning to forget: Continual prediction with LSTM. Neural computation.2000;12(10):2451–71.

30. Srivastava N, Hinton GE, Krizhevsky A, Sutskever I, Salakhutdinov R. Dropout: a simple way to prevent neural networks from overfitting. Journal of Machine Learning Research. 2014;15(1):1929–58.

31. Snoek J, Larochelle H, Adams RP, editors. Practical bayesian optimization of machine learning algorithms. Advances in neural information processing systems; 2012.

32. BayesianOptimization. GitHub repository. 2016.

33. Friedman JH. Greedy function approximation: a gradient boosting machine. Annals of statistics. 2001:1189–232.

34. Friedman J, Hastie T, Tibshirani R. Additive logistic regression: a statistical view of boosting (with discussion and a rejoinder by the authors). The annals of statistics. 2000;28(2):337–407.

35. Ho TK. The random subspace method for constructing decision forests. IEEE transactions on pattern analysis and machine intelligence. 1998;20(8):832–44.

36. Wolpert DH. Stacked generalization. Neural networks. 1992;5(2):241–59.

37. Cameron AC, Windmeijer FA. An R-squared measure of goodness of fit for some common nonlinear regression models.Journal of Econometrics. 1997;77(2):329–42.

38. Domencich TA, McFadden D. Urban travel demand-a behavioral analysis. 1975.

39. Amirikian B, Georgopulos AP. Directional tuning profiles of motor cortical cells. Neuroscience research. 2000;36(1):73–9.

40. Paninski L, Shoham S, Fellows MR, Hatsopoulos NG, Donoghue JP. Superlinear population encoding of dynamic hand trajectory in primary motor cortex. The Journal of neuroscience. 2004;24(39):8551–61.

41. Weber AI, Pillow JW. Capturing the dynamical repertoire of single neurons with generalized linear models. arXiv preprint arXiv:160207389. 2016.

42. Lipton ZC, Kale DC, Elkan C, Wetzell R, Vikram S, McAuley J, et al. The Mythos of Model Interpretability. IEEE Spectrum.2016.

43. Lau B, Stanley GB, Dan Y. Computational subunits of visual cortical neurons revealed by artificial neural networks. Proceedings of the National Academy of Sciences. 2002;99(13):8974–9.

44. Prenger R, Wu MC-K, David SV, Gallant JL. Nonlinear V1 responses to natural scenes revealed by neural network analysis.Neural Networks. 2004;17(5):663–79.

45. McAuley J, Leskovec J, editors. Hidden factors and hidden topics: understanding rating dimensions with review text.Proceedings of the 7th ACM conference on Recommender systems; 2013: ACM.

46. Simonyan K, Vedaldi A, Zisserman A. Deep inside convolutional networks: Visualising image classification models and saliency maps. arXiv preprint arXiv:13126034. 2013.

47. Ribeiro MT, Singh S, Guestrin C, editors. Why should i trust you?: Explaining the predictions of any classifier. Proceedings of the 22nd ACM SIGKDD International Conference on Knowledge Discovery and Data Mining; 2016: ACM.

48. Strobl C, Boulesteix A-L, Kneib T, Augustin T, Zeileis A. Conditional variable importance for random forests. BMC bioinformatics. 2008;9(1):307.

49. Bell DA, Wang H. A formalism for relevance and its application in feature subset selection. Machine learning.2000;41(2):175–95.

50. Corbett EA, Perreault EJ, Körding KP. Decoding with limited neural data: a mixture of time-warped trajectory models for directional reaches. Journal of neural engineering. 2012;9(3):036002.

51. Feurer M, Klein A, Eggensperger K, Springenberg J, Blum M, Hutter F, editors. Efficient and robust automated machine learning. Advances in Neural Information Processing Systems; 2015.

52. Kotthoff L, Thornton C, Hoos HH, Hutter F, Leyton-Brown K. Auto-WEKA 2.0: Automatic model selection and hyperparameter optimization in WEKA. Journal of Machine Learning Research. 2016;17:1–5.

